# ChIP-MS reveals the local chromatin composition by label-free quantitative proteomics

**DOI:** 10.1101/2023.01.27.525999

**Authors:** Wai Khang Yong, Grishma Rane, Nurkaiyisah Zaal Anuar, Xiaoman Shao, Chai Yeen Goh, Vartika Khanchandani, Vivian L. S. Kuan, Maya Jeitany, H. Phillip Koeffler, Lih-Wen Deng, Takaomi Sanda, Dennis Kappei

**Author notes:** Correspondence should be addressed to Dennis Kappei.

## Abstract

Chromatin immunoprecipitation (ChIP) has been a cornerstone for epigenetic analyses over the last decades, but even coupled to sequencing approaches (ChIP-seq), it is ultimately limited to one protein at a time. In a complementary effort, we here combined ChIP with label-free quantitative (LFQ) mass spectrometry (ChIP-MS) to interrogate local chromatin compositions. We demonstrate the versatility of our approach at telomeres, with transcription factors, in tissue and by dCas9-driven locus-specific enrichment.

---

Chromatin biology has been extensively studied by chromatin immunoprecipitation (ChIP)^1^, which is easily coupled to downstream next-generation sequencing (NGS) to study the global distribution of the target protein across the genome *in vivo* (i.e. one protein to many loci)^2^. While it would be equally informative to know comprehensively the chromatin composition at individual loci (i.e. one locus to many proteins), a widespread application of simple and robust DNA-centric approaches is still lacking. Beyond *in vitro* reconstitution methods^3, 4^, there have been increasing attempts to purify individual chromatin loci, e.g. through nucleic acid hybridization^5, 6^ or ChIP-based approaches coupled to mass spectrometry (ChIP-MS)^7–11^. Furthermore, recent approaches have exploited the sequence specificity of nuclease-dead Cas9 (dCas9) to target specific chromatin loci by directly enriching for dCas9^12–15^ or through proximity ligation using biotin ligases^16–20^. Despite these growing efforts, a major challenge remains to reliably study the protein composition at specific gene loci due to insufficient enrichment and suboptimal identification rates even of known factors^21^. Moreover, many approaches require either large amounts of input material (up to 10^11^ cells) or highly specialized reagents and plasmid vectors, making these techniques labour-intensive and challenging for routine analysis.

Furthermore, we argue that the gold standard for successful locus-specific approaches should be the identification of protein factors that independently and directly bind to the same chromatin fragment without involving protein-protein interactions. Here, we revisited ChIP-MS to develop a straightforward and robust workflow to study chromatin-bound proteins both genome-wide and at specific loci. We selected human telomeres as a widely used proof-of-concept locus, given that multiple independent direct telomere-binding proteins are well described. Telomeres, the ends of linear chromosomes, consist of short tandem repeats that are constitutively bound by the six-protein complex shelterin, containing TERF1 (also known as TRF1), TERF2 (also known as TRF2), POT1, TINF2 (also known as TIN2), ACD (also known as TPP1) and TERF2IP (also known as RAP1). This complex functions to prevent loss of downstream coding information and fusion of chromosomal ends^22^. Additional proteins have been described to directly bind telomeres without any known protein-protein interaction with the shelterin complex, including NR2C2^5, 23^, NR2F2^5, 23^, HMBOX1 (also known as HOT1)^4, 5^, ZNF827^24^ and ZBTB48 (also known as TZAP)^25, 26^. This well-described set of proteins makes telomeres an ideal locus to benchmark successful optimisation of ChIP-MS approaches.

To test whether independent telomere-binding proteins can reciprocally enrich each other, we performed standard double cross-linking ChIP reactions using the reversible cross-linkers formaldehyde (FA) and dithiobis(succinimidyl propionate) (DSP) for TERF2 and ZBTB48 in U2OS cells. Both proteins enriched telomeric DNA proportionate to their known relative abundance at telomeres^25^ (Extended Data Fig. 1a and 1b). ZBTB48, but not TERF2, was also enriched on the MTFP1 promoter as previously described^25^ (Extended Data Fig. 1b). Subsequently, we analysed equivalent TERF2 and ZBTB48 ChIP reactions by label-free quantitative (LFQ) mass spectrometry using 300 μg of DNA sonicate (equivalent to ~50 million U2OS cells) per replicate (Fig. 1a). TERF2 ChIP-MS significantly enriched for the bait protein TERF2 and all members of the shelterin complex compared to the IgG negative control (Fig. 1b, Supplementary Table 1). In addition, many factors known to more transiently associate with TERF2, the shelterin complex or telomeres such as SLX4, DCLRE1B, AURKB, MRE11A, BLM and TOP3A^22^ were significantly enriched. Importantly, as a proof that our approach genuinely identifies proteins independently bound to the same chromatin fragments, we also enriched other known direct telomere-binders, namely NR2C2, NR2F2, HMBOX1, ZNF827 and ZBTB48. In return, the reciprocal ZBTB48 ChIP-MS experiment enriched for a similar set of proteins compared to the IgG control (Fig. 1c, Extended Data Fig. 1c and Supplementary Table 2) including all members of the shelterin complex and other direct telomere-binding proteins. We further explored the presence of post-translational modifications in our mass spectrometry data and not only recapitulated key residues on TERF2, such as phosphorylation of Ser365 known to regulate telomeric t-loop formation^27^, but also modifications on independent telomere-binders such as Ser19 and Ser46 on NR2C2 (Extended Data Fig. 1d and Supplementary Table 3). The latter indicates that at least some of the telomere-bound NR2C2 carries these modifications. Next, to test whether our label-free approach allows for a seamless extension to tissue samples, we performed Terf2 ChIP-MS using 300 μg of sonicated chromatin from mouse livers of 7-week-old wildtype mice. We confirmed the successful enrichment of all shelterin members as well as the CST complex members CTC1 and OBFC1 as an independent telomere-binding complex^28, 29^ (Fig. 1d and Supplementary Table 4). Similar results were obtained with mouse embryonic stem cells as a species-matched cell line reference (Extended Data Fig. 1e-g and Supplementary Table 5). Finally, to showcase the applicability to DNA-binding proteins in general, we performed ChIP-MS for the transcription factor MYB in Jurkat cells, in which MYB acts as part of an oncogenic core regulatory circuit (CRC)^30^. In addition to the bait protein MYB, related protein factors such as TAL1, EP300, RUNX1, CBFB and GATA3 were enriched compared to the IgG control (Fig. 1e and Supplementary Table 6). Overall, these data showcase a straightforward ChIP-MS workflow that requires reasonably low input amounts, bypasses the need for any labelling, and is versatilely applicable to DNA-binding proteins from cell culture to tissue samples.

**Fig. 1.**
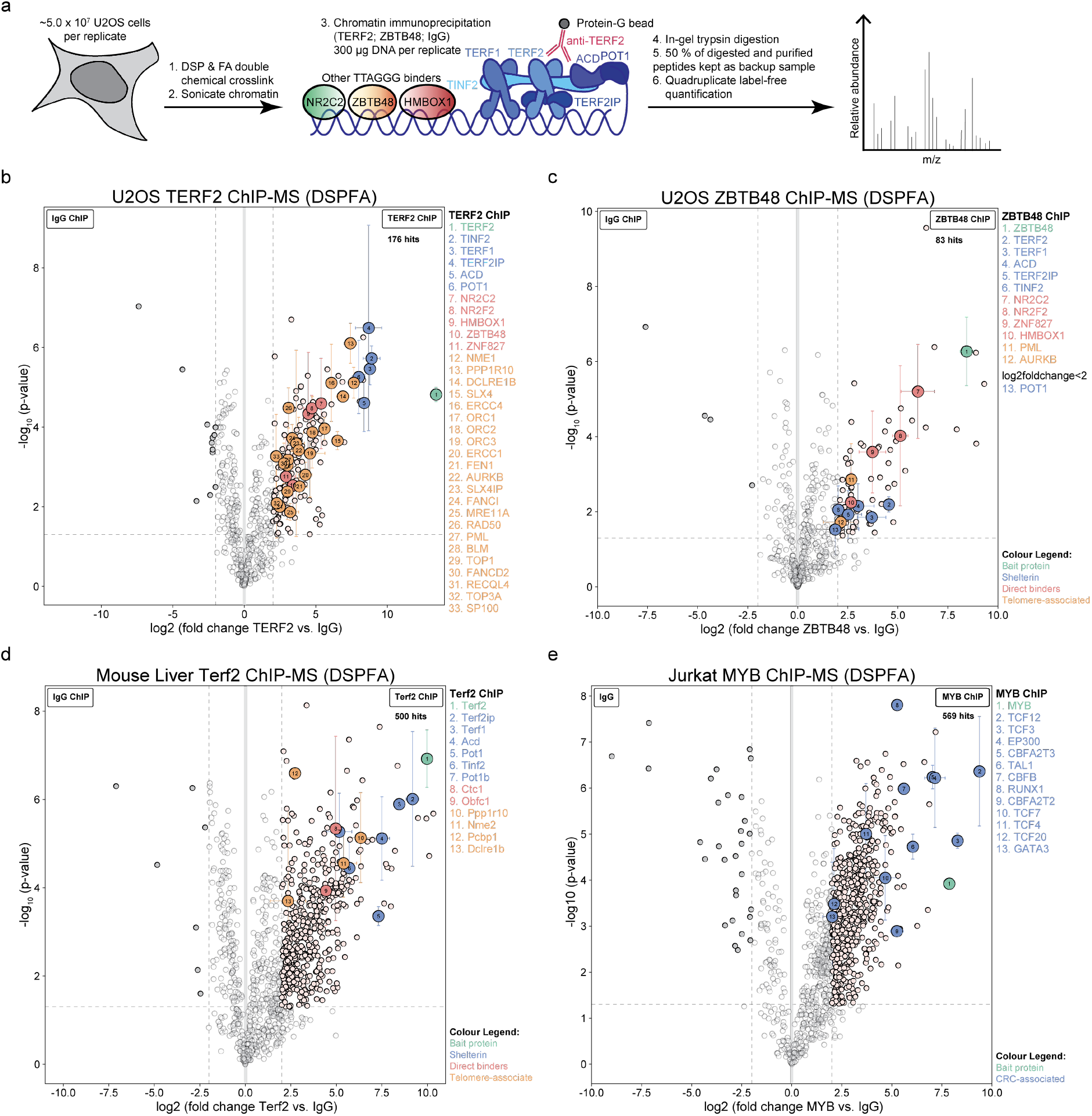
ChIP can be efficiently combined with label-free quantitative mass spectrometry. (a) Schematic of the label-free quantitative ChIP-MS workflow. (b-e) Volcano plots of ChIP-MS reactions for TERF2 (b) and ZBTB48 (c) in U2OS cells, Terf2 in mouse liver tissue (d) and MYB in Jurkat cells (e) (n = 4 biological replicates for panels b-e). Specifically enriched proteins are distinguished from background proteins by a two-dimensional cut-off of >4-fold enrichment and p<0.05. Two-dimensional error bars represent the standard deviation based on iterative imputation cycles during the label-free analysis to substitute missing values (e.g. no detection in the IgG control). (b-d) The bait proteins are marked in green, shelterin complex members are marked in blue, proteins known to independently bind to telomeres are marked in red and other known telomere-associated proteins are marked in orange. (e) The bait protein is marked in green, and CRC-associated proteins are marked in blue.

Similar to previous publications, our various ChIP-MS data enriched hundreds of proteins above our relatively stringent cut-offs. To test whether these results represent either the biological complexity of chromatin or whether cross-linked samples are more prone to false-positives, we repeated ZBTB48 ChIP-MS using previously established U2OS ZBTB48 KO clones^25^, allowing us to better account for antibody-specific background by avoiding the IgG control. Using each five U2OS WT and five ZBTB48 KO clones as biological replicates, the ChIP-MS comparison yielded a much smaller set of 27 proteins (Fig. 2a and Supplementary Table 7). While the shelterin proteins, independent direct telomere binders (NR2C2, NR2C1, NR2F2, HMBOX1 and ZNF827), and known telomere-associated proteins (ERCC4, TOP3A, BLM, PML and RAD50) were again significantly enriched, the majority of the remaining proteins – notably without prior functional association with telomeres – were here found within the background MS data (Extended Data Fig. 2a,b). Importantly, proteins common across all three telomeric U2OS ChIP-MS experiments (TRF2 vs. IgG, ZBTB48 vs. IgG. and ZBTB48 WT vs. KO) are mostly known telomere-associated proteins (Fig. 2b). These results may explain the generally low overlap between similar proteomics datasets, suggesting wide-spread inclusion of false-positives (Extended Data Fig. 2c,d). Our much shorter, rigorous hit list from the WT vs. KO comparison is in agreement with the challenging dynamic range of abundant chromatin proteins including histones^21^. This is further illustrated by the fact that independent chromatin fragment binders are reciprocally enriched with orders of magnitude lower signal intensities compared to protein-protein interactors (Fig. 2c). The stringent WT vs. KO comparison also allowed us to re-evaluate the requirement for double cross-linking for our ChIP-MS workflow. Indeed, using single FA cross-linking ZBTB48 ChIP-MS only yielded one shelterin member, TERF2, as well as NR2C/F nuclear receptors, highlighting the benefit to further stabilise interactions by DSP (Fig. 2d and Supplementary Table 8). However, constitutive KOs are only feasible for non-essential genes and might be undesirable for transcription factors such as MYB that affect a large set of target genes. Here, an alternative is rapidly inducible degron systems such as FKBP12-dTAG^31^. To showcase this, we generated a biallelic Jurkat FKBP12^F36V^-MYB knock-in clone (Extended Data Fig. 3a,b) and confirmed that MYB could be depleted by the addition of dTAG1^V^-1 within 1h (Extended Data Fig. 3c-e). In DMSO-treated cells, FKBP12^F36V^-MYB enriched a similar set of proteins as seen for MYB in Jurkat WT cells over the IgG control (Extended Data Fig. 3f,g and Supplementary Table 9). However, when performing MYB ChIP-MS comparing DMSO- and dTAG1^V^-1-treated cells only MYB and seven other proteins, including ZMYM4, were specifically enriched (Fig. 2e). In contrast, the plethora of proteins enriched in the IgG comparison (Fig. 1e and Extended Data Fig. 3f-h) were abundantly detected in the background (Extended Data Fig. 3i). Importantly, using the same MYB antibody in dTAG1^V^-1-treated cells against an IgG comparison still enriched the same large set of false-positive proteins, while MYB and ZMYM4 had strongly reduced enrichment ratios in agreement with little residual MYB protein remaining in these cells (Extended Data Fig. 3h). Overall, these data indicate that ChIP-MS is particularly prone to false-positives, likely due to the inherent cross-linking, and can be rigorously controlled via bait-specific loss-of-function conditions.

**Fig. 2.**
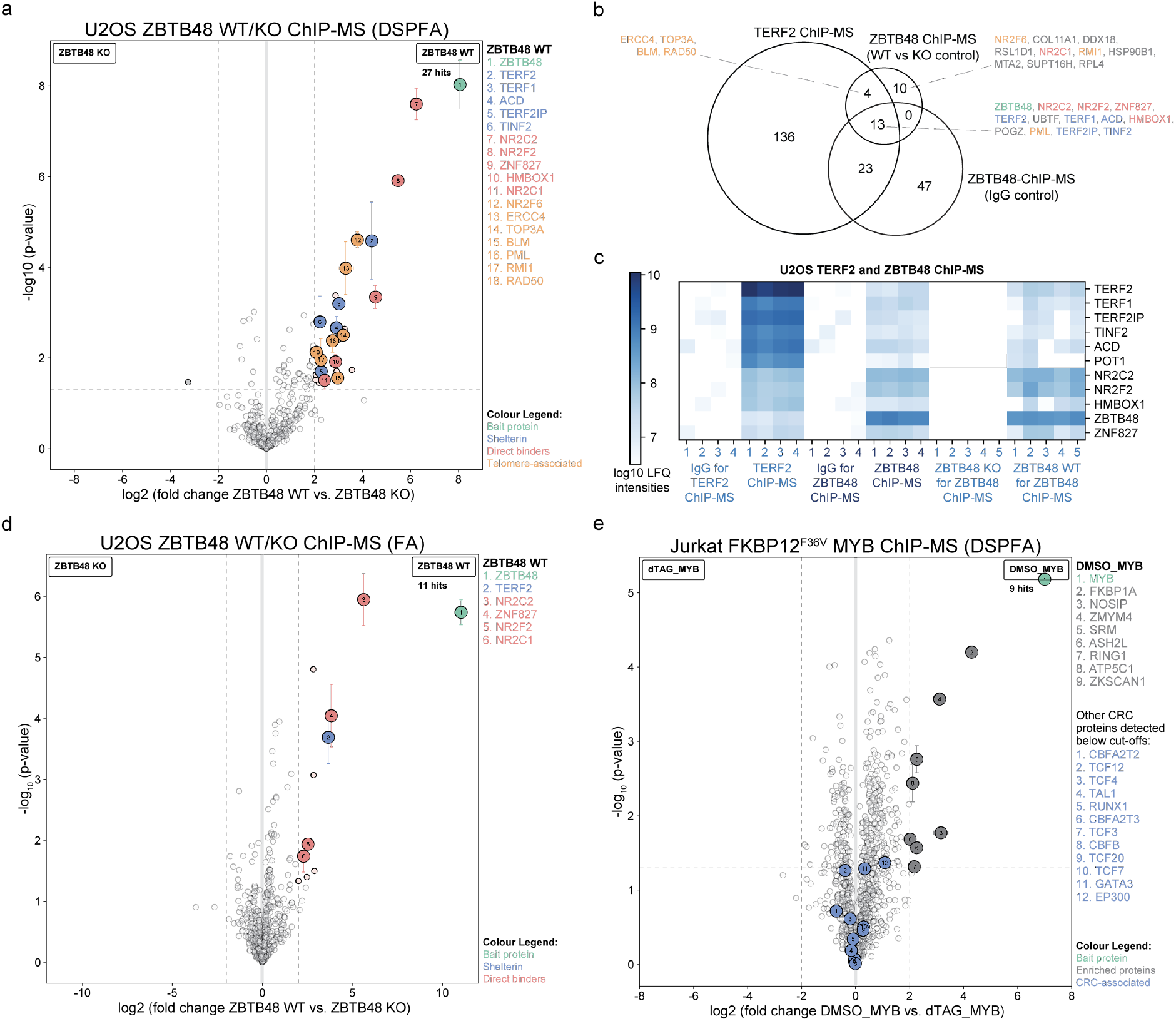
Loss-of-function controls are essential to exclude false-positives in ChIP-MS reactions. (a) ZBTB48 ChIP-MS reactions in five independent U2OS wildtype and ZBTB48 knock-out clones using double-crosslinking with FA and DSP (n = 5 biological replicates). (b) Venn diagram comparing telomere-associated proteins in TERF2 and ZBTB48 ChIP-MS reactions. (c) Heatmap visualization of log_10_ LFQ intensities for shelterin proteins and other direct telomere-binding proteins. (d) ZBTB48 ChIP-MS reaction in five independent U2OS wildtype and ZBTB48 knock-out clones using single-crosslinking with FA only (n = 5 biological replicates). (e) MYB ChIP-MS reaction using Jurkat FKBP12^F36V^-MYB knock-in cells upon 1 h treatment with either DMSO or 500 nM dTAG^V^-1 (n = 4 biological replicates). The volcano plots (a, d) use the same criteria as in Fig. 1b-d, while volcano plot (e) uses the criteria as in Fig. 1e.

While telomere-binding proteins predominantly enrich telomeric DNA, TERF2 and ZBTB48 have also been shown to bind and regulate gene bodies elsewhere in the genome^32, 33^. To address locus-specificity, we next incorporated dCas9 as a bait protein into our approach (Fig. 3a). To this end, we established a U2OS dCas9-GFP clone and subsequently transduced cells with sgRNAs targeting either telomeric DNA (sgTELO) or an unrelated control (sgGAL4). sgTELO- but not sgGAL4-expressing cells presented with a high degree of telomere localization (Fig. 3b) and enriched telomeric DNA in ChIP reactions (Fig. 3c-d). Furthermore, dCas9 ChIP followed by Western blot enriched TERF2 specifically in sgTELO samples (Fig. 3e). Finally, at the global level dCas9 ChIP-MS enriched all six shelterin members together with the telomere-associated proteins DCLRE1B and SLX4 as well as the independent telomere binders ZBTB48 and NR2C2 (Fig. 3f and Supplementary Table 10), demonstrating the capability of our label-free ChIP-MS approach for locus-specific applications.

**Fig. 3.**
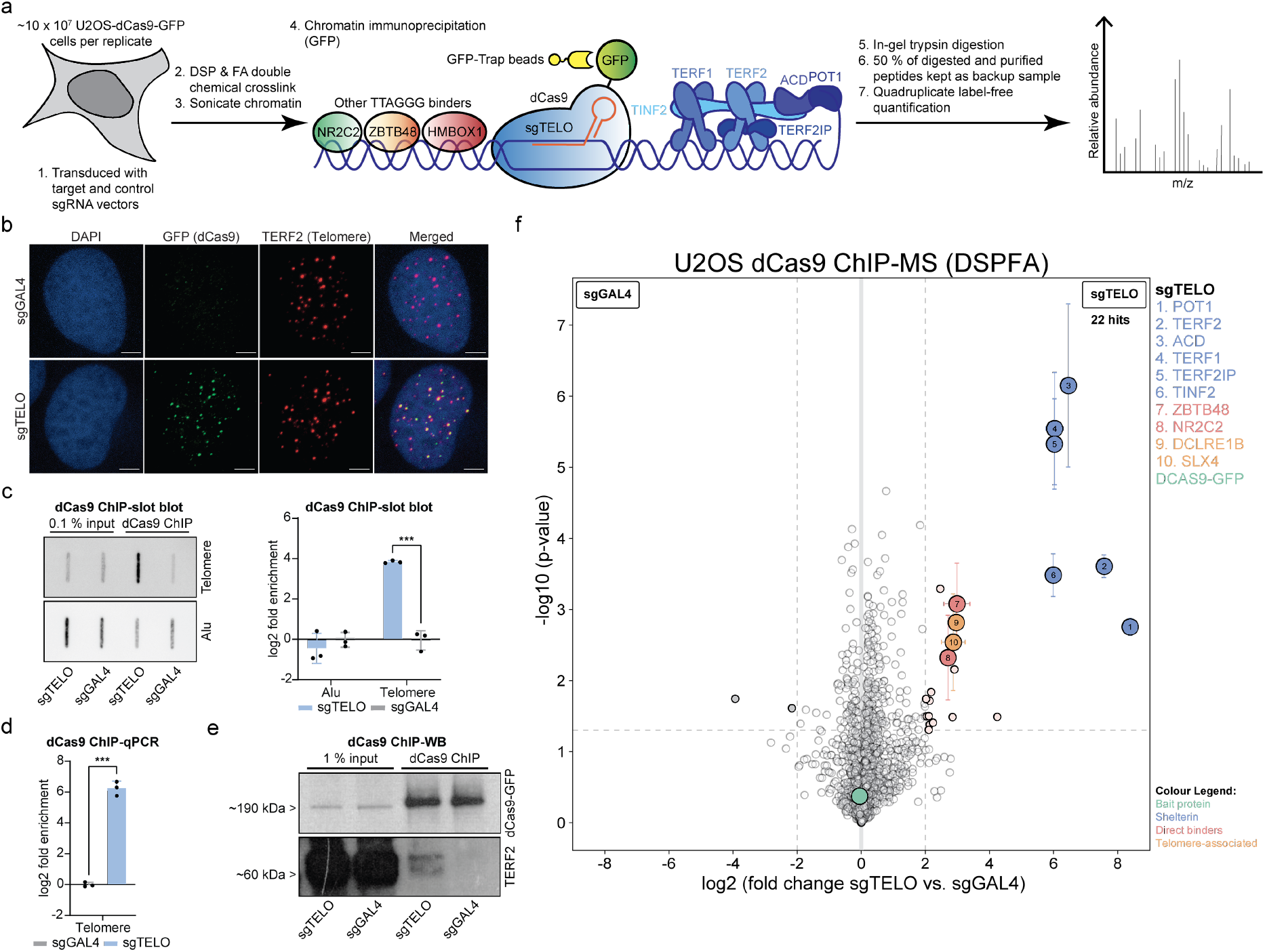
Locus-specific dCas9 ChIP-MS enriches known telomeric proteins. (a) Schematic of the label-free quantitative dCas9 ChIP-MS workflow. (b) Immunofluorescence microscopy of co-localization between dCas9-GFP (green) and TERF2 (red) in the presence of sgTELO or sgGAL4. Nuclei were counterstained with DAPI (blue). Scale bars represent 10 μm. (c) dCas9-GFP ChIP with either sgTELO or sgGAL4 analysed by slot blot with a probe for telomeric DNA or an Alu control (left). ChIP-slot blot quantification (right). (d) dCas9-GFP ChIP with either sgTELO or sgGAL4 analysed by qPCR. (e) dCas9-GFP ChIP with either sgTELO or sgGAL4 analysed by Western blot for dCas9-GFP and TERF2. (f) dCas9-GFP ChIP-MS reaction with either sgTELO or sgGAL4 (n = 4 biological replicates). The volcano plot uses the same criteria as in Fig. 1b-d.

Overall, we have here established a simple, robust ChIP-MS workflow based on comparably low input quantities^21^ that can be applied to any DNA-binding protein and in a locus-specific manner using dCas9. The choice of label-free quantitation provides a particularly low hurdle for laboratories without extensive mass spectrometry expertise and is readily compatible with tissue samples. In return, using a standard double cross-linking approach and widely used reagents for dCas9-GFP enrichment provides an easily applicable template for the myriad of labs with existing ChIP expertise. Critically, the complete enrichment of the shelterin complex in our dCas9 ChIP-MS example provides convincing proof-of-concept evidence that locus-specific ChIP-MS approaches are feasible despite well-reasoned concerns about possible steric hindrance^21^. Future generations of even more sensitive mass spectrometers and improvements in the experimental workflow, e.g. a combination with ChIP-SICAP^11^ or an adaptation of CUT&RUN^34^, are likely to push these boundaries further.

## Methods

### Data availability

The mass spectrometry data have been deposited to the ProteomeXchange Consortium via the PRIDE^35^ partner repository with the dataset identifiers PXD039627 and PXD039701.

## Acknowledgments

We are grateful to all members of the Kappei lab for advice and discussions. This research was supported by the National Research Foundation Singapore and the Singapore Ministry of Education under its Research Centres of Excellence initiative and an NMRC Open Fund Individual Research Grant (MOH-OFIRG21jun-011). Salary support for WK was provided by a NGS postgraduate fellowship and a NUSMed Postdoctoral Fellowship [NUSMED/2020/PDF/02].

## Author contributions

WK and DK conceived the study and designed experiments. WK performed experiments with help from GR, NZA, XS, CYG, VK, VLSK and MJ. HPK, LWD and TS contributed to the research supervision. WK and DK analysed the data. WK and DK wrote the manuscript with input from all authors.

## Materials and Methods

### Cell culture

Human U2OS and HEK293T cells were cultivated in Dulbecco’s modified Eagle’s medium (DMEM; Gibco) containing 4.5 g/l glucose, 4 mM glutamine and 1 mM sodium pyruvate, and human Jurkat cells were cultivated in Roswell Park Memorial Institute (RPMI) 1640 medium (Gibco). DMEM and RPMI media were supplemented with 10 % (v/v) foetal bovine serum (FBS; Gibco), 100 U/ml penicillin and 100 μg/ml streptomycin (Gibco). Mouse R1/E embryonic stem cells were cultivated in DMEM supplemented with 15 % (v/v) embryonic stem-cell FBS (Gibco), 10 mM MEK inhibitor PD0325901 (Selleckchem), 10 mM GSK-3 inhibitor CHIR-99021 (Selleckchem), 1000 U/ml mouse recombinant leukemia inhibitory factor (LIF; Gibco), 100 U/ml penicillin and 100 μg/ml streptomycin, 1 % (v/v) non-essential amino acids solution 100X (Gibco), and 0.1 % (v/v) 2-mercaptoethanol 1000X (Gibco). Mouse ES cells were provided with fresh medium every 48 h. All cells were cultured in a humidified incubator at 37 °C and 5 % CO_2_.

### Plasmids

The pLV-EF1a-dCas9-GFP-IRES-Neo vector was generated by cloning the dCas9-GFP sequence from vector pSLQ1658-dCas9-GFP into vector pLV-EF1a-IRES-Neo using EcoRI and BamHI. pSLQ1658-dCas9-GFP and pSLQ1651-sgTelomere(F+E) were gifts from Bo Huang & Stanley Qi^36^ (Addgene #51023 and #51024). pSLQ1651-sgRNA(F+E)-sgGal4 was a gift from Jian Xu^14^ (Addgene plasmid #100549). pLV-EF1a-IRES-Neo was a gift from Tobias Meyer^37^ (Addgene #85139). pRSV-Rev, pMDLg/pRRE and pMD2.G were gifts from Didier Trono^38^ (Addgene #12253, #12251 and #12259). The sgRNA vectors were cloned using pSLQ1651-sgTelomere(F+E) as the base plasmid as previously described^36^. pCRIS-PITChv2-Puro-dTAG (BRD4) was a gift from James Bradner & Behnam Nabet^31^ (Addgene #91793). pX330A-1×2 and pX330S-2-PITCh were gifts from Takashi Yamamoto^39, 40^ (Addgene #58766 and #63670).

### Generation of U2OS-dCas9-GFP clone and lentiviral transduction

Each lentiviral transfer vector (0.5 μg) was co-transfected with pMDLg/pRRE and pRSV-Rev (0.25 μg each), and envelope vector pMD2.G (0.25 μg) into 3 × 10^5^ HEK293T packaging cells in a 6-well plate. Plasmids were diluted in 50 μl Opti-MEM (Gibco) and mixed vigorously with 2.5 μg polyethylenimine (Polysciences) diluted in 50 μl Opti-MEM. Reaction mixes were incubated at room temperature for 20 min before transfection. Wells were washed after 18 h and fresh medium was replenished for lentivirus production. Lentiviral supernatants were collected 48 hours after transfection and filtered with 0.22 μm syringe filters, supplemented with 10 mM HEPES and 8 μg/mL polybrene (Sigma Aldrich), and used for transduction of 1.5 × 10^5^ U2OS cells in 6 well plates. U2OS cells were transduced with lentivirus for dCas9-GFP-IRES-Neomycin and selected with 400 μg/ml neomycin (Gibco) for at least 5 days. U2OS-dCas9-GFP cells were sorted into single cells with a FACSAria cell sorter (BD Biosciences) and screened by immunofluorescence following a subsequent lentiviral transduction with sgTELO. The single cell clone exhibiting the highest frequency of dCas9-GFP localization to the telomeres in the presence of sgTELO was selected and used for dCas9 ChIP-MS experiments. Cells derived from the single cell clone were transduced with either sgTELO or sgGAL4 as control and selected with 0.8 μg/ml puromycin (Gibco) for at least 2 days. Cells were continuously cultured in the presence of neomycin and/or puromycin.

### Generation of biallelic Jurkat FKBP12^F36V^-MYB knock-in clone

A Puro^R^-P2A-2xHA-FKBP12^F36V^-linker cassette was amplified from pCRIS-PITChv2-Puro-dTAG (BRD4) to include MYB microhomology sequences and a MluI restriction site (Supplementary Table 11). The cassette and pCRIS-PITChv2-Puro-dTAG (BRD4) backbone were digested with MluI-HF (NEB) and assembled into pCRIS-PITChv2-Puro-dTAG (MYB) with T4 DNA ligase (NEB). MYB N-terminal-specific sgRNA (GCTCCCCGTTACCTGTGCCG) was cloned into the pX330A-1×2 vector digested with BbsI-HF (NEB), and the PITCh sgRNA sequence from pX330S-2-PITCh was inserted with Golden Gate assembly (NEB) to generate pX330A-nMYB/PITCh. Jurkat cells were transiently transfected with the Neon Transfection System (Thermo Scientific). For each 100 μl electroporation tip, 2 μg pCRIS-PITChv2-Puro-dTAG (MYB) and 4 μg pX330A-nMYB/PITCh were transfected into 1 × 10^6^ cells at 1350 V, 10 ms, 3 pulses. Transfected cells were cultured for 2 days before selection with 1 μg/ml puromycin (Sigma) for at least 4 days. Subsequently, cells were split into single cells and cultured in 96-well plates. Cell pellets from single clones were lysed in 1× Cell Lysis Buffer (Cell Signaling Technology), and lysates were stored at −80 °C until use. Confirmation of homozygous FKBP12^F36V^ knock-in was validated by Western blot analysis. In addition, the knock-in alleles were further confirmed by RT-PCR combined with Sanger sequencing: RNA was extracted from the homozygous clone with NucleoSpin RNA Mini kit for RNA purification (Macherey-Nagel) and reverse transcribed into cDNA with EvoScript Universal cDNA Master (Roche). The N-terminal sequence of the 2xHA-FKBP12F36V-MYB transcript was PCR amplified with AccuPrime Pfx DNA Polymerase (Thermo Scientific) and purified with the QIAquick PCR purification kit (Qiagen) according to the manufacturer’s instructions (Supplementary Table 11). Sanger sequencing was performed on the purified PCR product (Supplementary Table 11).

### MYB protein degradation using dTAG1^V^-1

Jurkat FKBP12^F36V^-MYB cells were treated with dTAG^V^-1 (Tocris) dissolved in DMSO as a solvent. Optimized parameters for degradation of MYB were 500 nM dTAG^V^-1 for 1 h.

### Chemical double crosslinking

Cells grown on 15 cm cell culture dishes were fixed at room temperature with 2 mM DSP (Thermo Scientific) for 45 min, washed twice with ice-cold PBS, followed by a second fixation using 1 % (v/v) FA (Pierce) for 20 min at room temperature. FA was quenched for 5 min by adding 1.5 ml 2.5 M Glycine-PBS. Fixed cells were washed twice with ice-cold PBS, scrapped (for adherent cells) in additional 0.5 ml PBS with cOmplete proteinase inhibitor (Roche) with a cell scraper and stored as dry cell pellets in aliquots at −80 °C until lysis. Jurkat cells were fixed at room temperature in 2 mM DSP for 25 min before the addition of FA to a final concentration of 1 % (v/v) for an additional 20 min (i.e. 2 mM DSP for 45 min, 1 % (v/v) FA for 20 min). Fixed cells were washed twice with ice-cold PBS and stored as dry cell pellets in aliquots in −80 °C until lysis. Mouse livers were freshly harvested from 6-7-week-old C57BL6/NTac wildtype mice – in compliance with ethical regulations of the Institutional Animal Care and Use Committee (IACUC) of the National University of Singapore – and were promptly processed on ice. Each liver was processed individually by dicing into approximately 2 mm^3^ bits with scalpels on a 6 cm dish. Next, 4 ml of ice-cold PBS with protease inhibitor was added to each dish of processed liver samples, and the tissue slurry was pipetted into a pre-chilled glass dounce homogenizer and mechanically lysed on ice with 10 strokes using pestle A. The processed liver samples were transferred to 5 ml tubes and centrifuged at 700 g at 4 °C for 5 min. Supernatants were then discarded, and the processed liver samples were first fixed in 4 ml DMEM containing 2 mM DSP for 45 min at room temperature, washed twice with ice-cold PBS, followed by a second fixation in 4 ml DMEM containing 1 % (v/v) FA for 20 min at room temperature and two additional washes with ice-cold PBS. Tubes were incubated on a rotator during fixation. Processed and fixed liver samples were snap-frozen in liquid nitrogen and kept at −80 °C until lysis.

### Sonication of fixed samples

Preparation of sonicates was performed as previously described^25^ with modifications. Thawed cells were lysed with Lysis Buffer 1 (LyB1: 140 mM NaCl, 50 mM Tris-HCl (pH 8.0), 250 mM sucrose, 1 mM EDTA (pH 8.0), 10 % (v/v) glycerol, 0.5 % (v/v) Igepal CA-630, 0.25 % (v/v) Triton X-100, 0.25 % (v/v) Tween 20, cOmplete protease inhibitor for 15 min on a rotating wheel at 4 °C. Cell pellets were collected by centrifugation at 1,000 g for 10 min at 4 °C, and supernatants were discarded. Cell pellets were washed with Lysis Buffer 2 (LyB2: 200 mM NaCl, 10 mM Tris-HCl (pH 8.0), 1 mM EDTA (pH 8.0), 0.5 mM EGTA (pH 8.0), cOmplete protease inhibitor) for 10 min on a rotating wheel at 4 °C. Cell pellets were again collected by centrifugation at 1,000 g for 10 min at 4 °C, and supernatants were discarded. For every ^~^25 μl lysed cell pellet, approximately equivalent to 1 × 10^7^ U2OS cells, 150 μl of Sonication Buffer (50 mM Tris-HCl (pH 8.0), 10 mM EDTA (pH 8.0), 10 % (v/v) SDS, cOmplete protease inhibitor) was used to resuspend the pellet into a homogenous suspension prior to sonication in an ice bath using an EpiShear Probe Sonicator (Active Motif) with the following parameters: U2OS and R1/E – 30 % amplitude, 6.25 min sonication time (25 cycles of 15 s ON/ 30 s OFF); Jurkat – 30 % amplitude, 8.75 min sonication time (35 cycles); mouse liver tissues – 30 % amplitude, 7.5 min sonication time (30 cycles). Debris and dirt were cleared by centrifugation at 20,000 g for 10 min at 4°C. As a quality control, 8 μl sonicates were treated with RNase A (Thermo Scientific) and Protease K (Roche), purified with the QIAquick PCR purification kit (Qiagen) and DNA concentrations were estimated using the Qubit dsDNA HS Assay Kit (Thermo Scientific) according to manufacturers’ guidelines. Chromatin fragment sizes of 200-400 bp in length (lower – upper limits) were routinely obtained (determined by 1.5 % (w/v) agarose gel electrophoresis) and used for ChIP experiments.

### Chromatin Immunoprecipitation (ChIP)

DNA Lo-bind tubes (Eppendorf) were used throughout ChIP experiments. For each replicate, 25 μg DNA sonicate or approximately 4 × 10^6^ cells were mixed with 330 μl Protein Binding Buffer 1 (PBB1: **180 mM NaCl**, 50 mM Tris-HCl (pH 8.0), 0.25 % (v/v) Igepal CA-630, 1 mM DTT, 0.25 mM MgCl_2_, cOmplete protease inhibitor). 0.833 μg antibody was added to each ChIP reaction and incubated on a rotating wheel at 4 °C overnight: rabbit anti-TRF2 (Novus; #NB110-57130); rabbit anti-ZBTB48 (Sigma Aldrich; #HPA030417); rabbit anti-c-Myb phospho S11 (abcam; #ab45150), or rabbit IgG (ChromPure; #011-000-003). 12.5 μl Dynabead protein G magnetic beads (Thermo Scientific) were used for each ChIP reaction. For dCas9-GFP ChIP, 4 μl GFP-Trap magnetic agarose beads (Chromotek) were used in each reaction to enrich for dCas9-GFP overnight instead of antibodies. Magnetic beads were washed three times with PBB1 containing 10 μg/μl BSA at room temperature, by pipetting the beads in the solution 10 times and pelleted using a magnet bar. For the second wash, 3.33 μl 10 mg/ml sheared salmon sperm DNA (Thermo Scientific) per ChIP reaction was added, and the beads were washed using a rotating wheel for 10 min. On the following day, each ChIP reaction involving antibodies was transferred to a new tube containing washed Dynabead protein G and incubated at 4 °C for 2 h. **ChIP-Western blot (ChIP-WB)**: Magnetic beads were washed three times with Protein Binding Buffer 2 (PBB2: **150 mM NaCl**, 50 mM Tris-HCl (pH 8.0), 0.25 % (v/v) Igepal CA-630, 1 mM DTT, 0.25 mM MgCl_2_, cOmplete protease inhibitor). Beads were resuspended in 25 μl 2× Tris-Glycine Sample Buffer (Thermo Scientific). To reverse FA crosslinks, samples were incubated at 95 °C for 10 min. Denatured samples were stored at −80 °C until use. **ChIP-qPCR**: Magnetic beads were washed seven times with PBB2, and a final wash with ice-cold TE buffer. DNA was eluted twice using 100 μl Elution Buffer (1 % (w/v) SDS, 0.1 M NaHCO_3_) at 65 °C. Elutes were collected in clean DNA LoBind tubes (Eppendorf) and 8 μl 5 M NaCl was added to each sample, followed by overnight incubation at 65 °C. Input samples for qPCR analysis were treated similarly as the eluates. On the following day, samples were treated with RNase A and Proteinase K, purified with the QIAquick PCR purification kit (Qiagen) prior to qPCR analysis using the QuantiNova SYBR Green PCR Kit (Qiagen) on either QuantStudio 3 or 5 Real-Time PCR Systems (Applied Biosystems). See Supplementary Table 11 for a list of ChIP-qPCR primers and conditions used.

### Telomere slot blot analysis

Samples used for ChIP-qPCR were separately used for telomere slot blot analysis. ChIP samples were denatured at 80 °C for 10 min and immediately chilled on ice until samples were cooled to room temperature. The required volumes of samples were diluted to 200 μl with 2× saline-sodium citrate (SSC) Buffer (0.30 M NaCl, 0.03 M sodium citrate; pH 7) and set aside until use. A 48-well slot blotter (SCIE-PLAS) was assembled with a 12.8 cm × 4.4 cm positively charged nylon membrane. Wells were loaded and drained using a vacuum, first with 200 μl distilled water and then with 200 μl 2× SSC Buffer, prior to subsequent loading of diluted DNA samples into the required wells. A vacuum was applied to bind DNA samples onto the membrane. The DNA-bound membrane was first placed onto a piece of Whatman paper pre-soaked with Denaturing Solution (1.5 M NaCl, 0.5 M NaOH) at room temperature for 10 min, followed by placing the membrane on another piece of Whatman paper pre-soaked with Neutralizing Solution (3 M NaCl, 0.5 M Tris-HCl; pH 7.5) at room temperature for 10 min. Thereafter, bound DNA was fixed onto the blotting membrane by UV using 120 MJ before washing the membrane twice with 2× SSC Buffer. Probe hybridization and chemiluminescence detection was performed with the TeloTTAGGG Telomere Length Assay (Roche) according to the manufacturer’s recommendations. Bound DNA was probed with digoxigenin (DIG)-labelled telomere probes or DIG-labelled Alu control probes at 1:5,000 dilution.

### Western blot analysis

Cell lysates or samples denatured in 2× Laemmli Buffer (Sigma-Aldrich) or 4× LDS Sample Buffer (Thermo Scientific) were loaded on NuPAGE 4–12 % Bis-Tris Mini Protein gels (Thermo Scientific) in a XCell SureLock Mini-Cell electrophoresis system (Thermo Scientific) and resolved using 1× NuPAGE MOPS SDS Running Buffer (Thermo Scientific) at 170 V for 60 min. PageRuler Plus (Thermo Scientific) and MagicMark XP (Thermo Scientific) ladders were used routinely as standards. Methanol-activated PVDF membranes were used for both wet and semi-dry transfers with 1× Transfer Buffer (25 mM Tris, 192 mM glycine, 0.02 % (v/v) SDS, 10 % (v/v) methanol), in either a XCell II Blot Module (Thermo Scientific) at 35 V for 90 min or a Trans-Blot SD Semi-Dry Transfer Cell (Bio-Rad) at 60 mA per blot for 60 min, respectively. Membranes were subsequently blocked in 5 % (w/v) milk diluted in PBS containing 0.1 % (v/v) Tween 20 (PBS-T) at room temperature for 1 h. Membranes were incubated with primary antibodies overnight in 5 % (w/v) PBS-T, washed three times with PBS-T for 5 min each, incubated with secondary antibodies at room temperature for 1 h, and washed three times with PBS-T again. Protein bands were visualized with either Pierce ECL Western Blotting Substrate (Thermo Scientific) or ECL Select Western Blotting Detection Reagent (GE) using a ChemiDoc MP Imaging System (Bio-Rad). Primary antibodies: rabbit anti-TRF2 1:2,000 (Novus; #NB110-57130); rabbit anti-c-Myb phospho S11 1:500 (abcam; #ab45150); mouse anti-Myb 1:2,000 (Merck; #05-175); mouse anti-GFP 1:4,000 (Roche; #11814460001); mouse anti-tubulin 1:20,000 (MPI-CBG Antibody Facility); mouse anti-GAPDH 1:1,000 (Santa Cruz; #sc-47724). Secondary antibodies: goat anti-rabbit HRP conjugate 1:3,000 (Bio-Rad; #1706515); goat anti-mouse HRP conjugate 1:3,000 (Bio-Rad; #1706516).

### ChIP-mass spectrometry (ChIP-MS)

ChIP-MS was performed with quadruplicates for each experimental condition. The procedure is similar to ChIP-WB but optimized parameters were scaled up by a factor of 12. For each replicate 300 μg DNA sonicate (TERF2, ZBTB48, MYB), approximately 5 × 10^7^ cells, were mixed with 3960 μl PBB1, 10 μg antibodies and 150 μl Dynabead protein G in a 5 ml DNA LoBind tube (Eppendorf). For dCas9-GFP ChIP-MS, each reaction consisted of 600 μg DNA sonicate, approximately 10 × 10^7^ cells, 7920 μl PBB1 and 96 μl GFP-trap beads prepared in two 5 ml DNA LoBind tubes (Eppendorf). Protein samples were eluted by resuspending magnetic beads in 25 μl 2× Tris-Glycine Sample Buffer after the final three washes with PBB2. To reverse DSP crosslinks, 2 μl 1 M DTT were added, and samples were incubated at 37°C for 30 min, followed by 10 min at 95°C to reverse FA crosslinks. Denatured samples were stored in −80°C until sample preparation for proteomics analysis.

### Label-free quantitative proteomics

Denatured ChIP-MS elutes were separated by SDS-PAGE gel electrophoresis using NuPAGE 12 % Bis-Tris Mini Protein gels (Thermo Scientific) at 160 V for 20-30 min. Each ChIP-MS sample was prepared as two or four fractions for in-gel trypsin digestion. Briefly, gel fractions were cut approximately into 2 by 2 by 1 mm pieces, destained twice using Destaining Solution (50 % (v/v) ethanol, 25 mM ammonium bicarbonate) and dehydrated with acetonitrile. Gel pieces were dried using a speed vacuum concentrator (Eppendorf) and subsequently treated with Reducing Solution (50 mM DTT, 25 mM ammonium bicarbonate) at 56 °C for 1 h followed by Alkylating Solution (55 mM iodoacetamide, 25 mM ammonium bicarbonate) at room temperature for 45 min in the dark. Gel pieces were washed once with 50 mM ammonium bicarbonate and twice with acetonitrile, dried using a speed vacuum concentrator, and incubated with sequencing grade trypsin (Promega) in 50 mM ammonium bicarbonate at 37 °C overnight. Digested peptides were extracted from the gel pieces on the following day through two incubations each using an alternating cycle of Extraction Buffer (3 % (v/v) trifluoroacetic acid, 30 % (v/v) acetonitrile, 25 mM ammonium bicarbonate) and acetonitrile for 20 min and 10 min, respectively, followed by a final incubation in acetonitrile for 10 min. All supernatants were collected and combined. Extracted peptides in solution were concentrated using a speed vacuum concentrator for 2 h, and desalted with home-made stage-tips containing C-18 resins (Empore). Stage tips were stored at 4°C prior elution of peptides for analysis using an EASY-nLC 1200 Liquid Chromatograph (Thermo Scientific) coupled to either a Q Exactive HF (Thermo Scientific) or timsTOF fleX (Bruker) mass spectrometer.

#### Q Exactive HF

Peptides were separated on a C-18-reversed phase column (25 cm length, 75 μm inner diameter; New Objective) which was packed in-house with ReproSil-Pur 120 C18-AQ 1.9 μm resin (Dr Maisch). The column was mounted on an Easy Flex Nano Source and maintained at 40°C by a column oven (Sonation). A 105-min gradient from 2 to 40 % (v/v) acetonitrile in 0.1 % (v/v) formic acid at a flow of 225 nl/min was used. The spray voltage was set to 2.2 kV. The Q Exactive HF was operated with a TOP20 MS/MS spectra acquisition method per MS full scan, conducted with 60,000 at a maximum injected time of 20 ms and MS/MS scans with 15,000 resolution at a maximum injection time of 75 ms.

#### timsTOF fleX

Peptides were separated on an Aurora series column (25 cm length, 75 μm inner diameter, C-18 1.7 μm; IonOpticks) with an integrated captive spray emitter. The column was mounted on a captive spray ionisation source and temperature controlled by a column oven (Sonation) at 50°C. A 105-min gradient from 2 to 40 % (v/v) acetonitrile in 0.1 % (v/v) formic acid at a flow of 400 nl/min was used. The spray voltage was set to 1.65 kV. The timsTOF fleX was operated with data-dependent acquisition (DDA) in PASEF mode with 10 PASEF ramps per topN acquisition cycle (cycle time 1.17s) and a target intensity of 10,000. Singly charged precursor ions were excluded based on their position in the m/z-ion mobility plane and precursor ions that reached the target intensity were dynamically excluded for 24 seconds.

Raw files from both mass spectrometers were processed with MaxQuant^41^ version 2.0.1.0 with preset standard settings for label-free quantitation using at least 2 LFQ ratio counts (except dCas9 ChIP-MS data which was analysed with at least 1 LFQ ratio count). Carbamidomethylation was set as fixed modification while methionine oxidation and protein N-acetylation were considered as variable modifications. Searches were performed against the human (UP000005640; v20210307) or mouse (UP000000589; v20210307) Uniprot databases. Search results were filtered with a false discovery rate of 0.01. Known contaminants, proteins groups only identified by site, and reverse hits of the MaxQuant results were removed and only proteins with LFQ intensities were kept. As a default, protein entries with at least four non-zero values in either condition of each pair-wise comparison were included in the analyses. For the ZBTB48 WT/KO ChIP-MS experiments, involving five WT and five KO clones, at least three non-zero values were required. Missing LFQ intensity values were dealt with by randomly imputing a value within the lowest 5 % of LFQ intensities in the same set of MaxQuant-processed data using a normal distribution with the means of three iterations.

### Immunofluorescence

Immunofluorescence stainings were performed on glass coverslips in 12 well-plates. Cells were fixed with 10 % formalin solution (Sigma Aldrich) for 10 min at room temperature and washed twice with PBS + 30 mM glycine. Cells were then permeabilized with PBS + 0.5% Triton X-100 for 5 min at 4°C, followed by two washes with PBS + 30 mM glycine. Cells were then incubated in blocking buffer (PBS + 0.2 % (w/v) fish skin gelatin (Sigma Aldrich) for 15 min at room temperature. Primary antibodies were diluted in blocking buffer and incubated with the cells for 1 h at room temperature. The following primary antibodies were used: mouse anti-GFP 1:500 (Roche; #11814460001), rabbit anti-TRF2 1:750 (Novus; #NB110-57130). Coverslips were washed three times with blocking buffer for 5 min each, and incubated with secondary antibodies diluted in blocking buffer for 1 h at room temperature. The following secondary antibodies were used: donkey anti-mouse AlexaFluor 488 1:1,000 (Thermo Scientific; #A21202); donkey anti-rabbit AlexaFluor 647 1:1,000. (Thermo Scientific; #A32795). Coverslips were again washed three times with blocking buffer for 5 min each and once with PBS, and mounted on glass slides with ProLong Gold Antifade Mountant with DAPI (Thermo Scientific). Imaging for co-localization was conducted on a LSM 880 with Airyscan confocal microscope (Zeiss) with a 100x/1.4 oil objective.

### Data analysis

Statistical analysis was performed with GraphPad Prism 9. Each dataset was tested for normality to determine the appropriate use of parametric or non-parametric tests. Statistical analyses as well as number of replicates were described in each figure legend. Statistical significance was assigned as followed: p-value < 0.05: *; p-value < 0.01: **; p-value < 0.001: ***. For volcano plots, p-values were obtained by performing a standard independent 2 sample t-test using the scipy.stats module in python.

## Extended Data Figures

**Extended Data Fig. 1.**
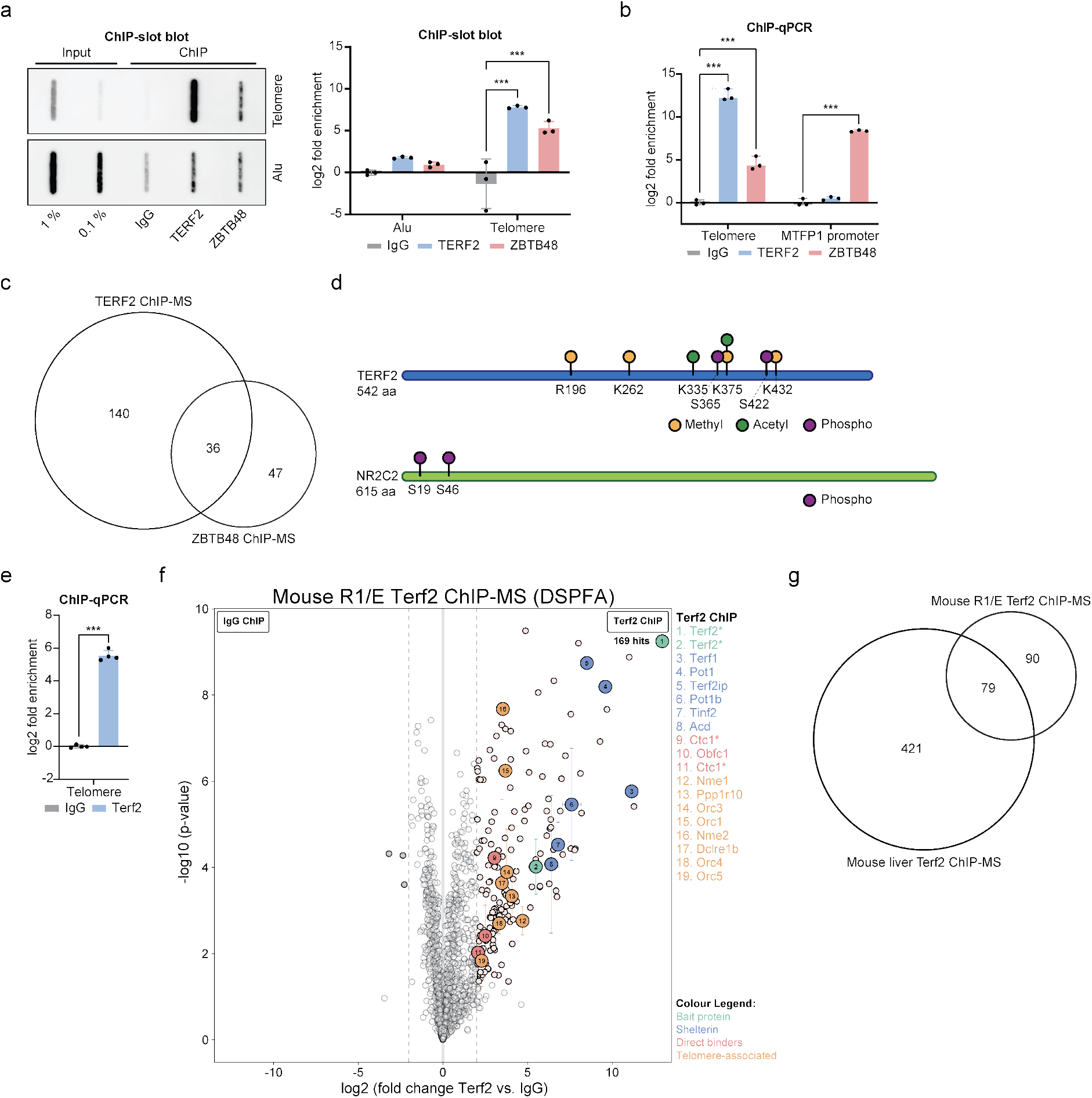
ChIP validations, analysis of PTMs and Terf2 ChIP-MS in mouse ES cells. (a) TERF2 and ZBTB48 ChIP reactions from U2OS cells analysed by slot blot with a probe for telomeric DNA or an Alu control (left). ChIP slot blot quantification (right). (b) TERF2 and ZBTB48 ChIP reactions from U2OS cells analysed by qPCR for telomeric DNA and the MTFP1 promoter. (c) Venn diagram comparing telomere-associated proteins in TERF2 and ZBTB48 ChIP-MS reactions using IgG as a negative control (Fig. 1b,c). (d) Schematic of identified post-translational modifications on TERF2 and NR2C2 in TERF2 and/or ZBTB48 ChIP-MS reactions (Fig. 1b,c). (e) Terf2 ChIP reactions from R/1E mouse embryonic stem cells analysed by qPCR for telomeric DNA. (f) Terf2 ChIP-MS reactions from R1/E mouse embryonic stem cells. The volcano plots use the same criteria as in Fig. 1b-d. Asterisks (*) denote proteins for which different isoforms, differentiated by unique peptides, were individually identified. (g) Venn diagram comparing telomere-associated proteins in Terf2 ChIP-MS reactions from R1/E mouse embryonic stem cells and mouse liver samples (Fig. 1d).

**Extended Data Fig. 2.**
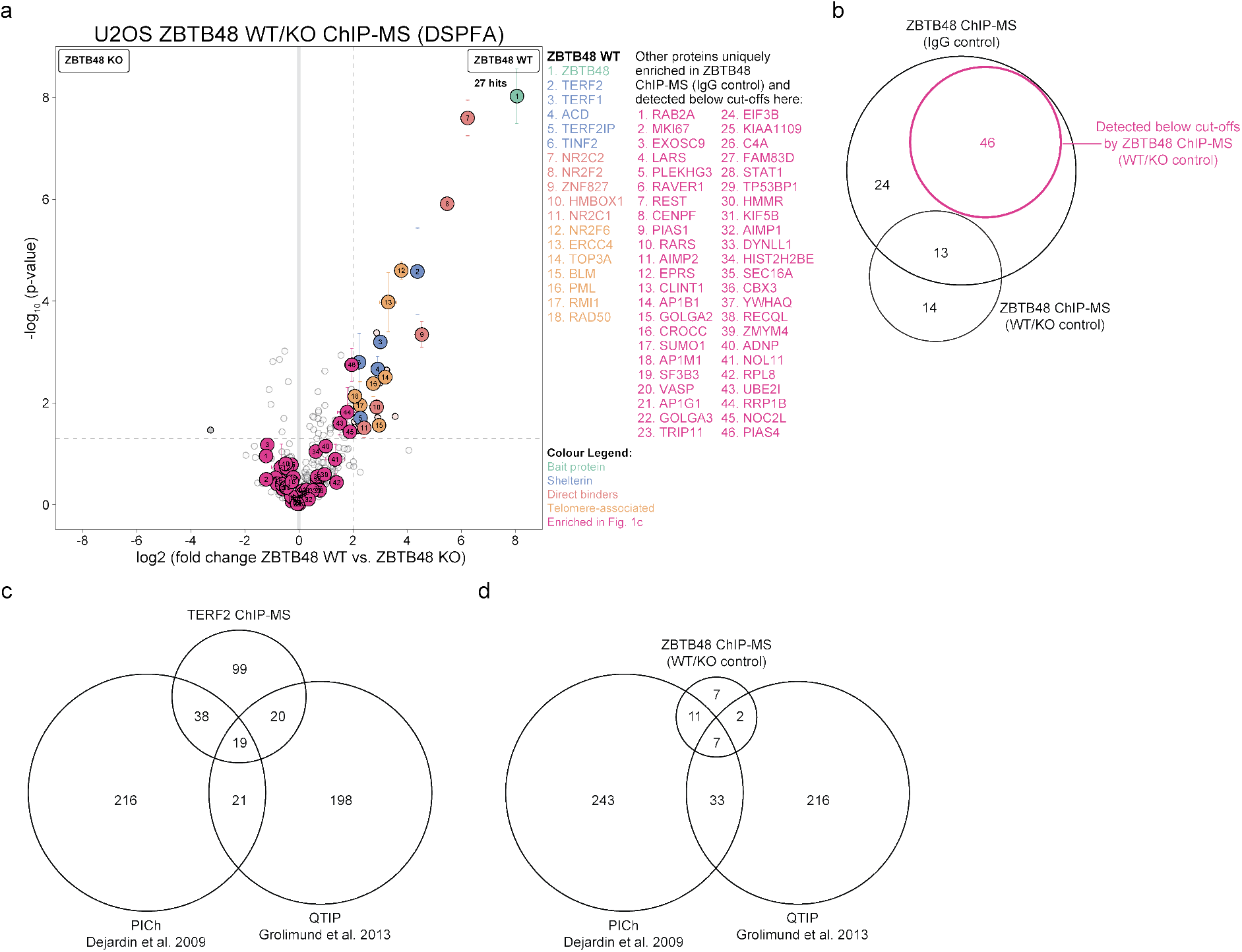
Knock-out controls exclude false-positives in telomeric ChIP-MS data. (a) ZBTB48 ChIP-MS reaction in five independent U2OS wildtype and ZBTB48 knock-out clones using double-crosslinking with FA and DSP (n = 5 biological replicates). Data as in Fig. 2a with the addition of protein hits above the fold enrichment and p-value cut-offs in the ZBTB48 vs. IgG ChIP-MS comparison in Fig. 1c highlighted in pink. (b) Venn diagram comparing telomere-associated proteins in ZBTB48 ChIP-MS reactions using either IgG as a negative control or ZBTB48 KO cells (while using the ZBTB48 antibody). The number of protein hits from the IgG comparison that are detected below the cut-offs in the WT vs. KO comparison is highlighted in pink. (c-d) Venn diagram comparing proteins enriched in the TERF2 (c) or ZBTB48 (d) ChIP-MS reactions with previously reported datasets using either PICh^5^ or QTIP^8^.

**Extended Data Fig. 3.**
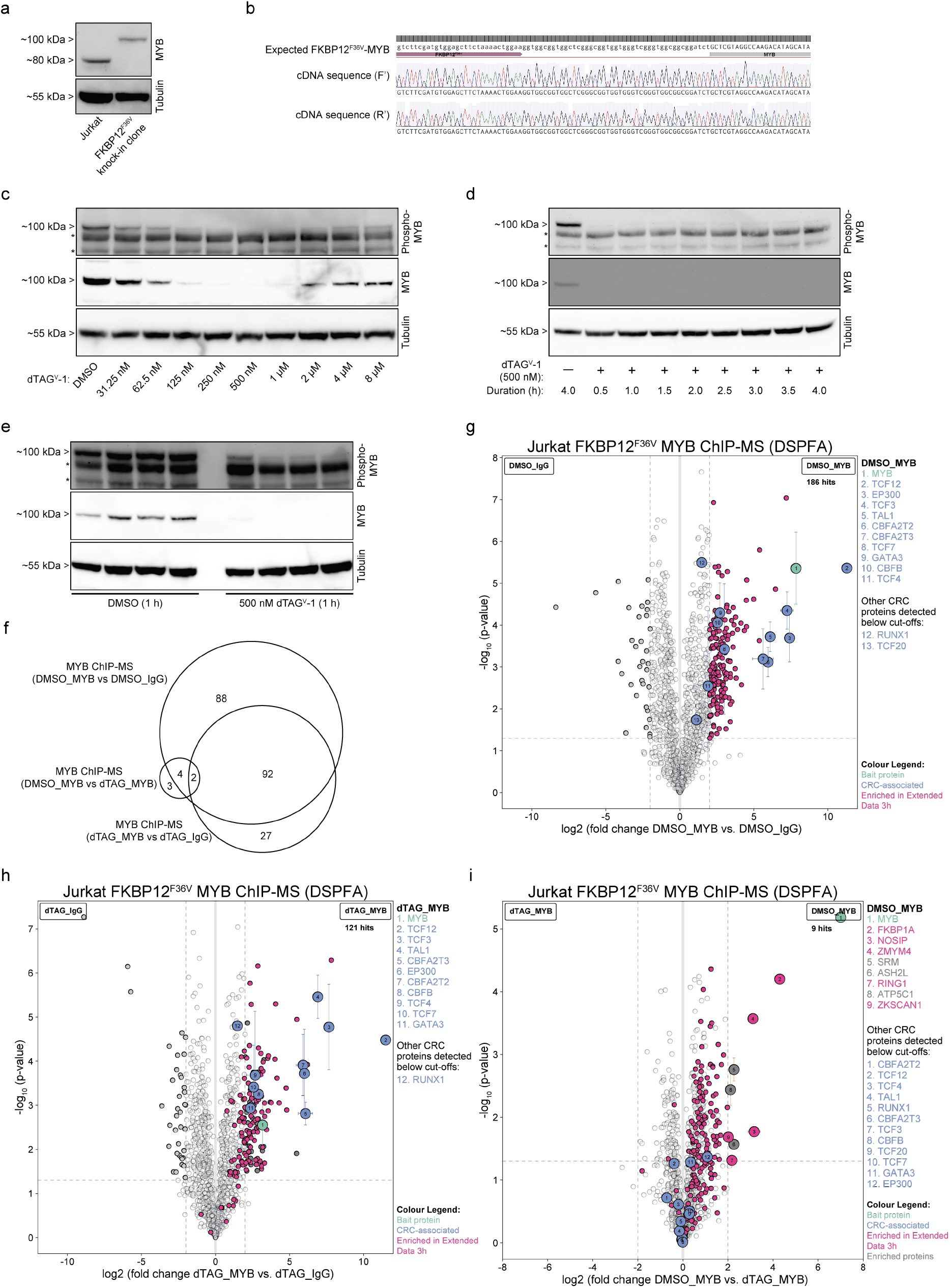
The dTAG-FKBP1 degron system excludes false-positives in MYB ChIP-MS data. (a) Western blot image of homozygous FKBP12^F36V^ knock-in clone showing a size shift of the MYB protein and a lack of wild-type MYB expression relative to parental Jurkat cells. Tubulin served as loading control. (b) Sanger sequencing validation of cDNA showing in-frame FKBP12^F36V^ insertion with no mutations. (c) dTAG^V^-1 dilution series to determine the optimal concentration. Jurkat cells were incubated for 1h with dTAG^V^-1 and protein extracts were anlysed by Western blot for MYB. Please note that the MYB phospho S11 antibody recognises unspecific bands (*) in addition to MYB (top band). Tubulin served as loading control. (d) Time course series to determine the optimal dTAG^V^-1 incubation time. Jurkat cells were incubated with 500 nM dTAG^V^-1. (e) DMSO vs. 500 nM dTAG^V^-1 treatment for 1h in Jurkat FKBP12^F36V^-MYB cells used as replicates for the ChIP-MS experiments (n = 4 biological replicates). (f) Venn diagram comparing enriched proteins in panels g-i. (g-i) MYB ChIP-MS reaction in Jurkat FKBP12^F36V^-MYB cells. (g) DMSO treated cells were used in comparison between MYB and an IgG control equivalent to the data of Jurkat WT cells in Fig. 2e. (h) dTAG^V^-1 treated cells were used in comparison between MYB and an IgG control upon 1 h incubation with 500 nM dTAG^V^-1. The volcano plots use the same criteria as in Fig. 1e. (i) DMSO vs. dTAG^V^-1 treated cells were used in MYB ChIP-MS reactions as in Fig. 2e with the addition of protein hits above the fold enrichment and p-value cut-offs in the MYB vs. IgG ChIP-MS comparison in Fig. 2e highlighted in pink.

## Supplementary Tables

**Supplementary Table 1 | U2OS TERF2 vs. IgG ChIP-MS data**

**Supplementary Table 2 | U2OS ZBTB48 vs. IgG ChIP-MS data**

**Supplementary Table 3 | U2OS TERF2/ZBTB48 vs. IgG ChIP-MS PTM data**

**Supplementary Table 4 | Mouse liver Terf2 vs. IgG ChIP-MS data**

**Supplementary Table 5 | R1/E mouse embryonic stem cells Terf2 vs. IgG ChIP-MS data**

**Supplementary Table 6 | Jurkat MYB vs. IgG ChIP-MS data**

**Supplementary Table 7 | U2OS ZBTB48 WT vs. KO ChIP-MS data using DSP-FA double crosslinking**

**Supplementary Table 8 | U2OS ZBTB48 WT vs. KO ChIP-MS data using FA single crosslinking**

**Supplementary Table 9 | Jurkat FKBP12^F36V^-MYB ChIP-MS data**

**Supplementary Table 10 | U2OS dCas9-GFP sgTELO vs. sgGAL4 ChIP-MS data**

**Supplementary Table 11 | Primer list for cloning, Sanger sequencing and ChIP-qPCR**

## Notes

### Competing Interest Statement

The authors have declared no competing interest.

